# Traditional uses and pharmacological activities of *Tetracera alnifolia* (Wild) Drake

**DOI:** 10.1101/2024.02.07.579282

**Authors:** A.K. Camara, E.S. Baldé, M.S.T. Diallo, M.K. Camara, T.V. Bah, M. Condé, A. Soumah, K. Kamano, I. Tietjen, A.M. Baldé

**Affiliations:** Faculty of Health Sciences and Techniques, Gamal Abdel Nasser University of Conakry. BP 1147, Guinea); Institute for Research and Development of Medicinal and Food Plants of Guinea-Dubréka, BP 6411, Guinea; National Institute of Public Health, Ministry of Health and Public Hygiene, Conakry, BP 6623, Guinea; The Wistar Institute, Philadelphia, Pennsylvania, USA

**Author notes:** **Corresponding author: Elhadj Saidou Baldé** Institute for Research and Development of Medicinal and Food Plants of Guinea-Dubréka, BP 6411, Guinea.

**Keywords:** *Tetracera alnifolia*, Ethnomedical Investigations, Biological Activity, SARS-CoV-2, Protozoa

## Abstract

**Ethnopharmacological relevance:** *Tetracera alnifolia* (Wild) Drake, is well used in traditional Guinean medicine for the treatment of infectious skin diseases. The present aim was to contribute to the valorization of *Tetracera alnifolia* leaves, focused on ethnomedical, biological and phytochemical investigations.

**Materials and methods:** we conducted an ethnomedical survey across several markets of the city of Conakry to identify 39 healers. Chloroform, methanol, dichloromethane, and aqueous extracts were tested for activities against protozoa, bacteria, fungi, HIV, and SARS-CoV-2.

**Results:** the traditional healers indicated that *T. alnifolia* is used in the treatment of more than 15 pathologies including *Fassa* (marasmus/malnutrition), *Soukhou kouyé* (white discharge in women), and *Tèmou bankhi* (sexual weakness in men). Leaves were the most used part. The modes of preparation included decoction and powder. Data from biological activities identicatied good activities of the methanolic extract against *Leishmania infantum* (MIC = 8.11 μg / ml) and a moderate activity on *Trypanosoma brucei* (MIC = 28.15 μg / ml) and *Staphylococcus aureus* (MIC = 29.91 μg / ml), while dichloromethane extracts acted on live SARS-CoV-2 replication with up to 53.4% inhibition at 50 μg/mL.

**Conclusion:** these results explain at least in part the traditional use of *T. alnifolia*

## Introduction

Before the discovery of antibiotics, infectious and parasitic diseases were the leading cause of death worldwide. They are now responsible for about 8% of deaths in developed countries but 50% of deaths in low-income countries (Murray et al., 2022). Within the context of increased poverty and malnutrition, combined with limited hygiene availability and access to high education, infectious and protozoan diseases affect millions of people every year in low-income countries and currently represent the primary cause of mortality in the tropical zone (Santos et al., 2020).

In Guinea, according to data from the Ministry of Health, eight neglected tropical diseases (onchocerciasis, lymphatic filariasis, trachoma, schistosomiasis, soil-transmitted helminths, leprosy, human African trypanosomiasis and Buruli ulcer), in addition to malaria, were considered a public health problem (Cherif et al., 2023). Also, more than 50% of the population lives in highly rural areas where access to conventional healthcare facilities is rare. Taking into account these conditions, rural people in Guinea rely strongly on traditional herbal medicine to manage their healthcare, including to treat infectious diseases. (A. Baldé et al., 2006; A. M. Baldé et al., 2015; A. Diallo et al., 2012; M. S. T. Diallo et al., 2019; Magassouba et al., 2007; Traore et al., 2013).

*Tetracera alnifolia* Willd is a perennial, evergreen big liana of the family Dilleniaceae which commonly grows in the forests and other warm regions of sub-Saharan Africa including Guinea. Several parts of *T. alnifolia* have been traditionally used for treating infectious diseases (including sexual transmitted disease), skin diseases, and malaria in Guinean traditional medicine. This study is part of the program of the Institute for Research and Development of Medicinal and Food Plants of Guinea (IRDPMAG), whose goal is to rationalize the integration of phytotherapy into our health systems as recommended by the WHO. In this context, based on the ethnomedical investigations conducted by IRDPMAG, *Tetracera alnifolia* was selected to confirm some of its traditional uses in Guinea as well as to survey its chemical and biological properties.

## 1. Materials and Methods

### 1.1. Ethnomedical survey

Our survey took place across several markets in Conakry, Guinea including Niger, Madina, Bonfi, Gbessia, Koloma, Enco 5, Taouyah, Kenien and Matoto markets. We opted for individual contact with healers in these markets. For consenting healers, we used the interactive method of interviewing the healers about their knowledge in the form of a semi-structured interviews following the outline of the survey questionnaire. Each questionnaire included demographic information such as sex, age, status but also description of diseases according to the healers, the mode of acquisition of the knowledge, and the modes of preparation and administration of the recipes. The study protocol was approuved by the IRDPMAG ethic committee.

### 1.2 Plant material

*T alnifolia* material was collected in the prefecture of Dubréka (Guinea). Samples were dried at room temperature. A voucher specimen (44HK470) was deposited in the herbarium of IRDPMAG.

### 1.3 Preparation of plant extracts

*T alnifolia* material was air dried and then ground in a mill. Powders (10 g) were submitted to maceration for 24 hours with 100 mL chloroform, 100 mL methanol and 150 mL of distilled water, separately. The chloroform and methanol extracts were filtered, pooled, evaporated *in vacuo* at < 40 °C in a rotary evaporator, coded (TaCHCl_3_; 20 mg) (TaCH_3_OH; 40 mg) respectively and stored at −80 °C until tested. The filtered aqueous extract was lyophilized to give aqueous extract (TaH_2_O; 50mg).

For SARS-CoV-2 test a part of aqueous extract (20 mg) was dissolved in water and treated by dichloromethane (100 mL). The dichloromethane fractions were filtered, pooled, evaporated *in vacuo* at < 40 °C in a rotary evaporator to give a dichloromethane extract (TaCH_2_Cl_2_; 3mg)

### 1.4 Biological testing

#### 1.4.1 Antibacterial and antifungal screening

The following strains of bacteria and fungi were used: *Escherichia coli* ATCC-8739, *Staphylococcus aureus* ATCC-6538 *Mycobacterium chelonae* and *Candida albicans* ATCC-10231. Estimation of the minimal inhibitory concentration (MIC) was carried out by the broth dilution method (M. A. Baldé et al., 2020; Cos et al., 2006). 100 µL of a bacterial or yeast suspension was aliquoted into 96-well plates and subsequently 100 µL of a twofold plant extract dilution was added to each well at desired concentrations. A control for normal microbial growth consisting of media without extract was added. Culture broths used were tryptic soy broth (TSB) for bacteria and Sabouraud (SAB) for fungi. The wells were inoculated with a microbial suspension of 10^5^ CFU/mL and then incubated for 24 hours. The inhibition of bacterial or yeast growth was evaluated by comparing wells with extracts to wells with normal microbial growth in the control wells without plant extracts. The minimal inhibitory concentration (MIC) was determined as the lowest concentration that completely inhibited macroscopic growth of bacteria or yeast. Samples with a MIC value of < 64 µg/mL were considered active. Ampicilin (Fluka), rifampicin (Fluka) and flucytosine (Sigma) were used as positive controls. These reference compounds are routinely used in the screening platform, and their activities were in the range that is usually observed.

#### 1.4.2 Anti-HIV assay

Anti-HIV screening against HIV-1 (strain IIIb) and HIV-2 (strain ROD) was carried out as reported before (Maregesi et al., 2010). Testing of anti-HIV activity used is based on evaluation of the inhibitory effect of a test agent against the cytopathic effects of the HIV-virus in a T-cell model using MT-4 cells. Azidothymidin was used as a positive control.

#### 1.4.3 Anti-SARS-CoV-2 assay

Vero-E6 cells were obtained from the American Tissue Culture Collection and cultured in D10+ medium consisting of Dulbecco’s Modified Eagle Medium, 10% fetal bovine serum (Gemini Bio Products, West Sacramento, USA), 100 U of penicillin/mL, and 100 μg of streptomycin/mL (Sigma Aldrich, St. Louis, USA) at 37 °C and 5% CO_2_. The following reagent was deposited by the Centers for Disease Control and Prevention and obtained through BEI Resources, NIAID, NIH: SARS-Related Coronavirus 2, Isolate USA-WA1/2020, NR-52281. To generate virus stocks, 3 x 10^6^ Vero-E6 were incubated in 15 mL of D10+ for 24 hours, replaced with fresh media, and incubated with virus at a multiplicity of infection of 0.001. Cells were incubated for 5-7 days until clear cytopathic effect was observed. Media was harvested and stored at -80 °C. To determine virus titers, cells were plated in 96-well format at 2 x 10^4^ cells/mL, incubated for 24 hours, and then washed and incubated in fresh media containing 5-fold serial dilutions of thawed virus aliquot for 4 days. Wells were then scored visually for the presence of cytopathic effect, and 50% tissue culture infectious dose TCID_50_ was calculated using the Reed-Muench method.

Antiviral screening was performed as described in Tietjen et al., 2021. Briefly, Vero-E6 cells were plated in D10+ medium at 2 x 10^4^ cells/mL in 96-well format, and extracts were added to cells at stated concentrations in triplicate. After 2 hours, cells were infected with 150x TCID_50_ of SARS-CoV-2 (USA-WA1/2020 variant). After 4 days incubation, cells were treated with resazurin to a final concentration of 20 μg/mL and incubated for an additional 4 hours. Cells were fixed with 4% paraformaldehyde for 30 minutes to inactivate virus, and cell viability was measured using a ClarioStar plate reader (BMG Labtech) (Tietjen et al., 2021). Remdesivir was included as a reference control. To determine effects of extracts on cell viability in the absence of infection, uninfected cells were cultured in parallel as described.

#### 1.4.4 Antitrypanosomal activity

Antitrypanosoma assays were performed as described by Baldé (E. S. Baldé et al., 2010).

##### Trypanosoma brucei

Briefly, Trypomastigotes of *T*.*brucei* Squib-427 strain (suramin-sensitive) were cultured at 37 °C and 5% CO_2_ in Hirumi-9 medium, supplemented with 10% fetal calf serum (FCS). Assays were performed by adding 1.5 x 10^4^ Trypomastigotes/well. After 72 hours incubation, parasite growth was assessed fluorometrically by adding resazurin for 24 hours at 37 °C. Fluorescence was measured using a Genios Tecan fluorimeter (excitation 530 nm, emission 590 nm). Suramin was included as a reference drug.

##### Trypanosoma cruzi

Tulahuen CL2 strain (nifurtimox-sensitive) was maintained on MRC-5 cells in minimal essential medium (MEM) supplemented with 20 mM L-Glutamine, 16.5 mM sodium hydrogen carbonate and FCS (5%) at 37 °C and 5% CO_2_. To determine *in vitro* Antitrypanosomal activity, 4 x 10^3^ MRC-5 cells and 4 x 10^4^ parasites were added to each well of test plate with compound. After incubation at 37 °C for 7 days, parasite growth was assessed by adding of beta-galactosidase substrate, chlorophenol red beta-D-galactopyranoside for 4 hours at 37 °C. The color reaction was read at 540 nm and absorbance values were expressed as a percentage of the blank controls. Nifurtimox was included as a reference drug.

#### 1.4.5 Antileishmanial activity

*Leishmania infantotum* amastigotes (MHOM/ET67) were collected from an infected donor hamster and used to infect primary peritoneal mouse macrophages as described (Corral et al., 2013). To determine *in vitro* Antileishmanial activity, 3 x 10^4^ macrophages were seeded in each well of a 96-well plate. After 48 hours outgrowth, 5 x10^4^ amastigotes/well were added and incubated for 2 hours at 37 °C. Pre-diluted compounds were subsequently added and the plates were further incubated for 120 hours at 37 °C and 5% CO_2_. Parasite burdens were determined microscopically after Giemsa staining and expressed as a percentage of the blank controls without compound. Pentostam was included as reference drugs.

#### 1.4.6 Antiplasmodial activity

Antiplasmodial activity evaluations were carried out as previously described (E. S. Baldé et al., 2010). The chloroquine-susceptible *P. falciparum* GHA-strain was used. Parasites were cultured in human erythrocytes A+ at 37 °C under a low oxygen atmosphere (3% O_2_, 4% CO_2_, and 93% N_2_) in a modular incubation chamber.The culture medium was RPMI-1640, supplemented with 10% human serum. 200 μL of infected human red blood cells suspension (1% parasitemia, 2% hematocrit) were added to each well of the plates with test compounds and incubated for 72 hours. After incubation, test plates were frozen at -20 °C. Parasite multiplication was measured by the Malstat method. 100 μL of Malstat reagent were transferred in a new plate and mixed with 20 µL of the haemolysed parasite suspension for 15 minutes at room temperature. After addition of 20 µL nitroblue tetrazolium (NBT)/ phenazine ethosulfate (PES) solution and 2 hours incubation in the dark, the absorbance was spectrophotometrically read at 655 nm using a Biorad 3550-UV microplate reader. Percentage growth inhibition was calculated compared to the negative blanks. Artesunate and chloroquine were included as reference drugs.

## 2. Results

### 2.1 Ethnomedical survey

The results of the ethnomedical surveys are shown in Tables 1 to 3 (see below).

**Table 1:** Sociodemographic data of healers.

| **Characteristics** | **Number of healers** | **Percentage (%)** |
| --- | --- | --- |
| **Age groups** |  |  |
| 30-40 | 9 | 23.1 |
| 41-50 | 13 | 33.3 |
| 51-60 | 11 | 28.2 |
| > 60 | 6 | 15.4 |
| **Total** | 39 | 100.0 |
| **Sex** |  |  |
| Male | 16 | 41.0 |
| Female | 23 | 59.0 |
| **Total** | 39 | 100.0 |
| **Mode of acquisition of knowledge** |  |  |
| **Mode of acquisition** |  | **Percentage (%)** |
| Family legacy | 33 | 84.6 |
| Learning | 4 | 10.3 |
| Former patients | 1 | 2.6 |
| Dreams | 1 | 2.6 |
| **Total** | 39 | 100.0 |

**Table 2:**
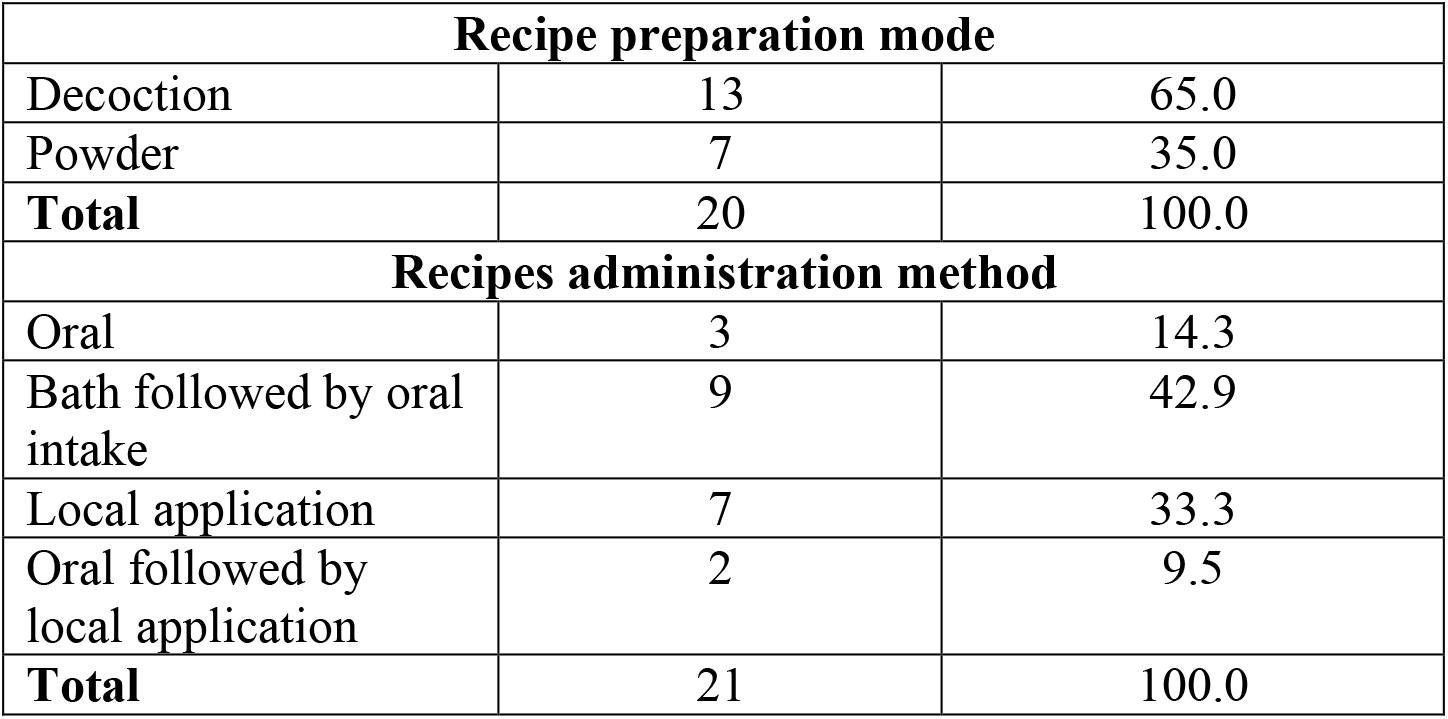
Methods of preparation and administration of recipes:

| **Recipe preparation mode** |  |  |
| --- | --- | --- |
| Decoction | 13 | 65.0 |
| Powder | 7 | 35.0 |
| **Total** | 20 | 100.0 |
| **Recipes administration method** |  |  |
| Oral | 3 | 14.3 |
| Bath followed by oral intake | 9 | 42.9 |
| Local application | 7 | 33.3 |
| Oral followed by local application | 2 | 9.5 |
| **Total** | 21 | 100.0 |

**Table 3:** Descriptions of diseases according to healers and ethnomedicals uses of *T. alnifolia*.

| **Vernacular name** | **Language** | **Symptoms treated** | **Approximate correlation** | **Number of quoted healers** |
| --- | --- | --- | --- | --- |
| *Makourou* | Maninka / Sosso | Warming of the belly and pain, pimples on the body, abdominal pain, stomach ache | Pyoderma | 4 |
| *Gnoni* | Maninka | Warming of the body, eyes and tongue are red and very painful | Varicella | 1 |
| *Wagna* | Maninka | Buttons, itching | Scabies | 5 |
| *Cassi* | Sosso | Itching, small pimples | Scabies | 2 |
| *Makourou yègbè* | Sosso | Fever and the body part is red | Pyoderma | 2 |
| *Cassi khounkhouri* | Sosso | Small pimples and itching | Scabies | 1 |
| *Soukhou kouyé* | Sosso | White fluid loss, pain on urination, itching, unpleasant odour in women. | Urogenital infections | 18 |
| *Fassa* | Maninka | Significant weight loss with the appearance of nerves on the skin in children. | Marasmus in children | 22 |
| *Konobori* | Maninka | Stools passed several times with fluid loss, sometimes with or without fever. | Diarrhea | 3 |
| *Temou Banki* | Sosso | Sex doesn't get up | Erectile dysfunction | 5 |
| *Mangué fakhè* | Sosso | Stomach ache, pain, constipation, anal itching, sometimes bleeding. | Hemorrhoids | 3 |
| *Danawali* | Sosso | Yellow tegument, fever, headache, chills, fatigue, vomiting | Malaria | 2 |
| *Faté fountou* | Sosso | Soft tissue swelling, sometimes a little painful | Heart failure | 2 |
| *Takoé* | Sosso | Whitish deposit on baby's tongue, pain, greenish stools. | Oral candidiasis | 2 |
| *Koundimi gbèlè* | Maninka | Severe head pain, difficulty opening eyes. | Sinusitis | 1 |
| *Sorom dimi* | Maninka | Pain in the pelvic region | Kidney disease | 1 |
| *Gnama* | Maninka | Lumbago with bleeding, nausea, headache, cramp in women. | Dysmenorrhea | 2 |
| *Fassa dimi* | Sosso | Pain, numbness, burning sensation in hands or feet | Neuropathy | 1 |

**Table 3:**
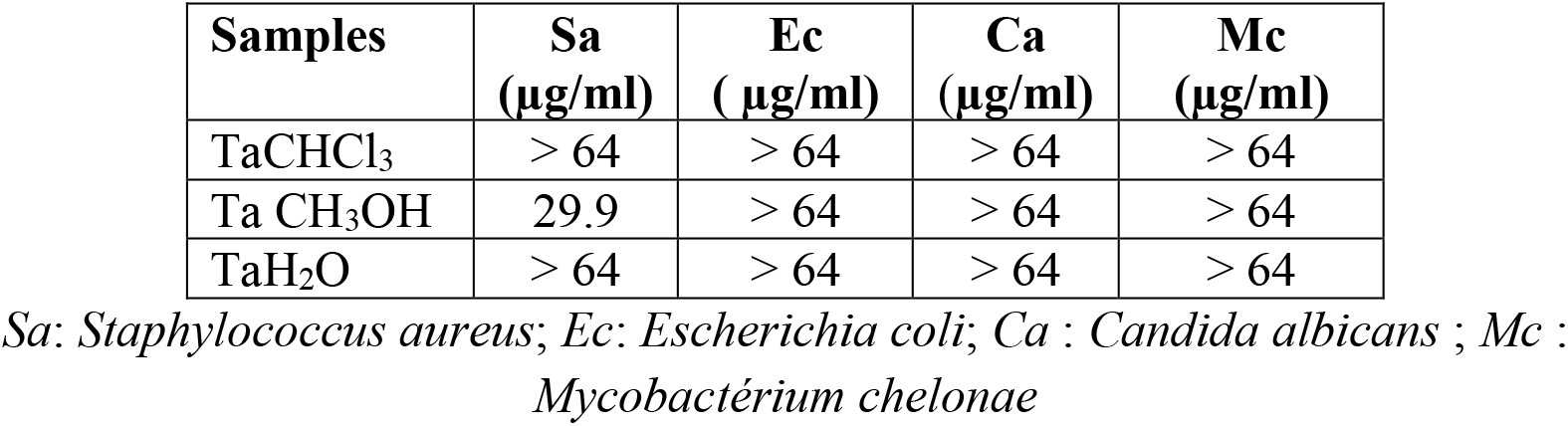
*in vitro* antibacterial and antifungal activities of extracts *T*.*alnifolia* (MIC in µg/mL).

From the ethnomedical survey, *Tetracera alnifolia* was also found to have numerous uses in Guinean traditional medicine (Table 3).

### 2.2. Biological activities

Three extracts (Chloroformic, methanolic and aqueous) were tested for antimicrobial activities. The antimicrobial effects of the extracts are summarized in Table 3. Aqueous and chloroformic extracts were not active against bacterial and fungal strains tested, while the methanolic extract was active against the *Staphylococcus aureus* strain (MIC = 29.9 µg/mL).

Chloroformic, methanolic and aqueous extracts were next tested for antiprotozoal activity as are summarized in Table 4. None of tested extracts were active against *Plasmodium falciparum* strain, while chloroformic extract was active against both trypanosma and leishmania strains.

**Table 4:**
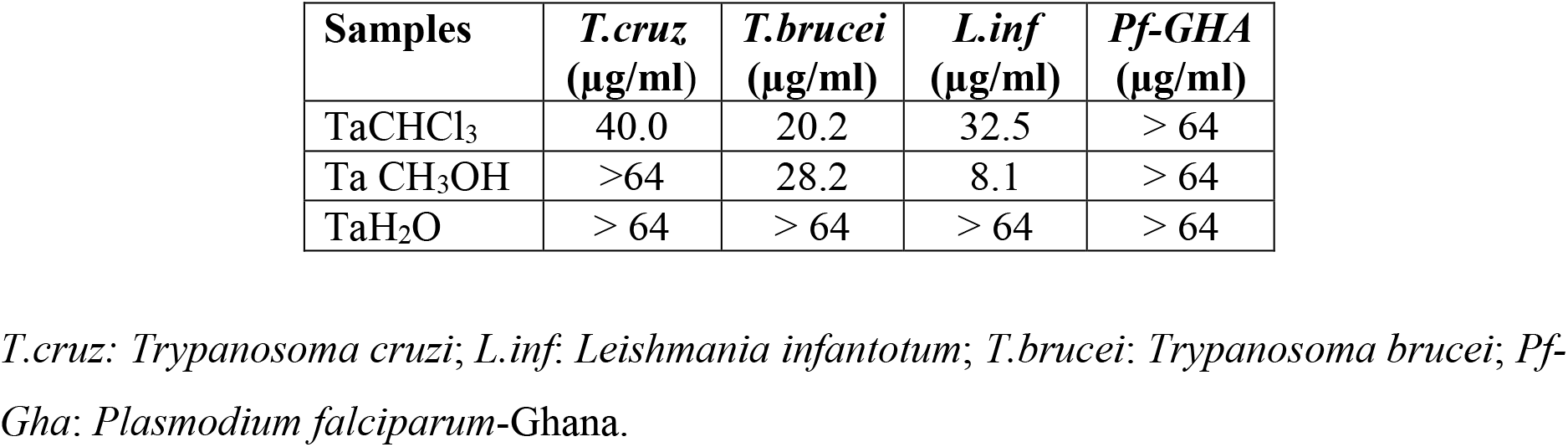
anti-trypanosomiasis, anti-leishmanial and anti-plasmodial activities *in vitro* extracts of *T*. *alnifolia* (MIC in μg/mL).

| Samples | *T.cruz*<br>(µg/ml) | *T.brucei*<br>(µg/ml) | *L.inf*<br>(µg/ml) | *Pf-GHA*<br>(µg/ml) |
| --- | --- | --- | --- | --- |
| TaCHCl<sub>3</sub> | 40.0 | 20.2 | 32.5 | > 64 |
| Ta CH<sub>3</sub>OH | >64 | 28.2 | 8.1 | > 64 |
| TaH<sub>2</sub>O | > 64 | > 64 | > 64 | > 64 |

*T.cruz*: *Trypanosoma cruzi*; *L.inf*: *Leishmania infantotum*; *T.brucei*: *Trypanosoma brucei*; *Pf-Gha*: *Plasmodium falciparum*-Ghana.

With regard to HIV, the chloroformic, methanolic and aqueous extracts showed MICs of 57.7, 62.7, and 52.6 µg/mL, respectively against HIV-1 IIIB strain, with less overall toxicity against in MT-4 cells (index of selectivity > 2). None of extracts tested was active against HIV-2 strain.

Chloroformic extract, methanolic and dichloromethane extracts were also tested for ability to inhibit live SARS-CoV-2 replication (parental Wuhan variant) in Vero-E6 cells (Figure 1). In this assay, chloroformic extract inhibited up to 31.4 ± 15.5% (mean ± s.e.m.) of virus replication relative to untreated, infected cells at 25 μg/mL, while extensive cell death was observed at 50 μg/mL. For dichloromethanic extract, up to 53.4 ± 5.4% of virus replication was inhibited at 50 μg/mL. In uninfected cells assessed in parallel, no toxicity was observed (i.e., > 90% viability relative to untreated cells) at any concentration up to 50 μg/mL (Figure 1), indicating good *in vivo* tolerability of these extracts. The additional toxicitity observed in infected cells treated with 50 μg/mL of chloroformic extract, relative to uninfected cells, is therefore likely to reflect an additive effect of both virus infection and high concentration of chloroformic extract.

**Figure 1:**
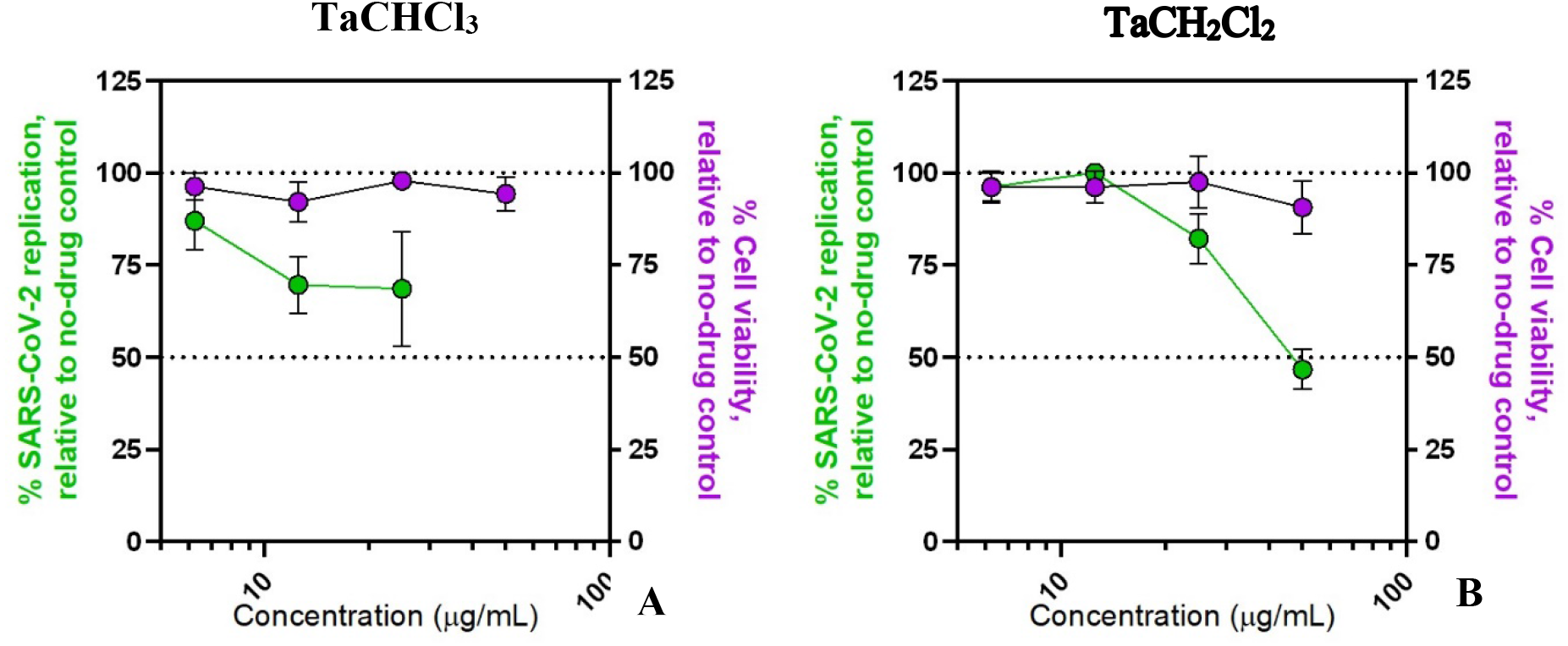
Dose response curves of chloromethanic extract A (left) and dichloromethanic extract B (right) on viral replication in SARS-CoV-2 USA-WA1/2020 variant-infected Vero-E6 cells (green) and cell viability in uninfected Vero-E6 cells (purple). Data are presented as percent SARS-CoV-2 replication and percent cell viability relative to uninfected or untreated cells, respectively, with 0% denoting viability of infected cells without treatment (for infection experiments) or media-only control (for cell viability experiments). Results show the mean ± s.e.m. of three independent experiments.

## 3. Discussion

Herbal medicine is a practice widely used by the population of Guinea for the management of several pathologies. This ethnomedical survey was conducted in some markets of the city of Conakry on the traditional use of *Tetracera alnifolia*. In total, we identified 39 healers, including 23 women (59.0%; Table 1) who participated in our survey. Indeed, this predominance of women is justified by the fact that women tend to be more interested in traditional medicine in this area. This result is somewhat different from that of Magassouba et al. (2007), who found a male predominance of healers (56.6%).

The age of healers was between 30 and 80 years old, and the most represented group was 41 to 50 years old with a frequency of 33.3% (13/39), followed by 51 to 60 years old with a frequency of 28.2% (11/39) (Table 1). This is in agreement with the previous results described in Guinea (A. Diallo et al., 2012; Magassouba et al., 2007). These results show that young people are less likely to be involved in the trades of traditional healers and indicates a need to promote traditional medicine among youth to prevent a gradual disappearance of this protected traditional knowledge. We also noted in our survey that 71.8% (28/39) of healers were not enumerated by the Ministry of Health compared to 28.2% (11/39) that were previously identified. This census deficit makes it difficult to organize and control the activities of healers, as well as to preserve and standardize their knowledge, and suggests that additional advocacy for healer organizations in Guinea is needed. The transmission of this traditional art is mainly inherited (84.7%, or 33/39) (Table 1). This result is similar to that reported by Traoré (Traore et al., 2013), who in an ethnobotanical survey on malaria in Guinea, reported a predominance of the acquisition of traditional art by inheritance of 50%, while acquisition by learning was much less common, i.e. 10.3% (4/39)

During our investigation, we identified six (6) pathologies mentioned by the healers of which the most cited were “*Wagna*” (5 quotes), “*Makourou*” (4 quotes), “*Cassi*” (2 quotes) and “*Makourou” yègbè* “(2 quotations) (Table 2). According to the descriptions of these pathologies by the healers, these affections could be of allergic origin (*Cassi khoun khouri, Wagna*). The most cited method of preparation was decoction (13 citations) followed by powder (7 citations) (Table 1). This result is similar to that of Diatta et al.; in their study carried out on the medicinal plants used against dermatoses in the Baïnounk pharmacopoeia of Djibonker, Senegal where the decocted ones were the most cited (Diatta et al., 2013). A decoction would be expected to collect the most active ingredients and mitigate or cancel the toxic effect of certain recipes (Zhang et al., 2018). Mainly, these forms of presentations, although easy to prepare, have certain disadvantages, including poor conservation, which results in the appearance of pathogenic molds and microorganisms in the recipes over time. We noticed in our investigation that *T. alnifolia* is also used in the treatment of other pathologies apart from dermatological conditions. These include “*Fassa*” (marasmus or malnutrition), “*Tèmou bankhi*” (sexual weakness in men) and “*Soukhou kouyé*” (white losses in women) (Table 3).

Three extracts of *T*.*alnifolia* were tested *in vitro* against microorganisms. Among the tested extracts, only the methanolic one presented a modest activity against *S. aureus*. Activity of methanolic extracts maybe the intermediary products between of chloroformic and aqueous extracts. None of these extracts was active against *Plasmodium falciparum*-Ghana. Previous investigations on biologically active plants have shown that crude extracts are usually more active than individual constituents of the plant (Rasoanaivo et al., 2011). However, both chloroformic and methanolic extracts of *T*.*alnifolia* moderately inhibited tested protozoa (*Trypanosoma* and *Leishmania*). The antiprotozoal that we observe here is in accordance with results reported by Traoré et *al*, (Traore et al., 2014). The lack of novel drugs for animal and human trypanosomiasis remains chronic and acute. Drugs resistance may perhaps best be countered by biologically active crude extracts from plants rather than the use of combinations of single chemical drugs known to be effective despite the risks of dual resistance (Cheuka et al., 2016) Available drugs for human trypanosomiasis now seem to be under production but have significant restrictions and are administrable with reasonable safety only under hospitalization (Kasozi et al., 2022)

Among the extracts tested in HIV, only the aqueous extract one presented a weak effect against HIV-1 IIIB strain. In contrast, two apolar extracts (Chloroform and dichloromethane) inhibited live SARS-CoV-2 replication at 12.5 to 50 μg/mL, depending on the extract, indicating the potential to identify natural product-based antivirals from this source.

## Conclusion

This study, whose purpose was to contribute to the enrichment of the ethnomedical, biological and phytochemical data of *Tetracera alnifolia*, made it possible to contact 39 healers who frequent markets in Conakry. The ethnomedical survey revealed that the plant is used in the traditional treatment of several diseases including skin infections, marasmus, white discharge in women and sexual weakness in men. Biotic investigations of the chloroform, methanol and aqueous extracts of the leaves showed good activity on protozoa including *Leishmania infantum* and Trypanosoma *brucei*. The methanolic extract showed moderate activity on *Staphylococcus aureus*, while aqueous and dichloromethane extracts acted on HIV-1 and SARS-CoV-2, respectively. Taken together, these studies indicate that *Tetracera alnifolia* extracts have wide traditional use in Guinea and a variety of anti-infective properties *in vitro*.

## Acknowledgements

We thank the Proteomics and Biochemistry of Proteins, University of Mons; Department of Organic and Biomedical Chemistry, NMR and Molecular Imaging Laboratory, University of Mons; Department of Pharmaceutical Sciences, University of Antwerp, Universiteits plein 1, B-2610 Antwerp, Belgium for the support, as well as the traditional practitioners who agreed to participate in their survey for their frank collaboration. Funding was provided by the Canadian Institutes for Health Research (CIHR PJT-153057) to I.T.

